# Towards a Brain-Computer Interface (BCI) for Improving Phonological Processing in Developmental Dyslexia: An Exploratory Study

**DOI:** 10.64898/2026.01.23.700941

**Authors:** Xuanci Zheng, João Araújo, Quentin Busson, Usha Goswami

## Abstract

Brain-computer interfaces (BCIs) have immense potential regarding the provision of therapies for disorders of development, but to date have typically been created for non-linguistic disorders such as ADHD (attention deficit hyperactivity disorder). Here we present a BCI designed to improve linguistic phonological processing during natural speech listening, aimed at participants with developmental dyslexia. Phonological ‘deficits’ are considered a core feature of dyslexia across languages and are present from infancy in those later diagnosed with dyslexia. Here a non-invasive EEG-BCI relying on auditory inputs and visual feedback was developed to optimise brain patterns related to phonological development in infants and children. The BCI focused on neural patterns identified using Temporal Sampling theory, which has found the theta/delta ratio during natural speech listening to be a key neural associate of impaired phonology. As a first step, the BCI was tested with adults. Adults with and without dyslexia played the BCI for 16 sessions, and received pre– and post-testing regarding phonological awareness and single word and nonword reading skills. Learning was assessed both online, as in most BCI studies, and offline, enabling removal of a greater number of potential EEG artifacts. Significant associations between offline BCI scores (a measure of BCI learning) and improvements in nonword reading were found for all participants who learned the BCI, both dyslexic and control. BCI learning also showed significant associations with improvements in real word reading and amplitude rise time discrimination. The data are interpreted with respect to Temporal Sampling theory.

## Introduction

Theories of developmental dyslexia attempt to provide a systematic causal framework for understanding this specific learning difficulty (e.g., Magnocellular theory, Stein & Walsh, 1997; Visual Attention Span theory, Valdois et al., 2004; Sluggish Attentional Shifting theory, Facoetti et al., 2010, Temporal Sampling theory, Goswami, 2011). A focus on *development* is absolutely critical to identifying core factor/s for effective remediation, accordingly here a brain-computer interface (BCI) is developed to address the phonological ‘core deficits’ that identify participants with dyslexia across languages (Stanovich, 1998). Importantly, in contrast to magnocellular and attention deficits (Goswami, 2022a for review), phonological deficits *pre-date* learning to read and are measurable in at-risk infants (Kalashnikova et al., 2019, 2020). The phonological ‘core deficit’ is operationalized using the cross-language sensory/neural Temporal Sampling theory (Goswami, 2011, 2015, 2018), which has identified key sensory and neural features of natural speech processing that are associated with impaired phonological development in infants and children (Goswami et al., 2002; Kalashnikova et al., 2018; Araújo et al., 2024; Attaheri et al., 2024). These sensory/neural features are thought to affect automatic speech processing from infancy onwards, resulting in the cognitive phonological impairments that are the hallmark of dyslexia in all languages (Ziegler & Goswami, 2005).

By Temporal Sampling theory, infants and children extract a phonological system (comprising phonological representations of the sound structures of individual words) via automatic sensory-motor learning of a hierarchy of amplitude modulations nested in the amplitude envelope of the speech signal (Goswami, 2022b). This phonological learning is an automatic outcome of the natural acquisition of spoken language. Accordingly, the phonological system emerges long before reading instruction commences. The BCI developed here focuses on phonological learning at the level of speech rhythm patterns, the key factor that governs infant language acquisition across all languages studied to date (Mehler et al., 1988; Nazzi et al., 1998). The phonological processing of speech rhythm patterns remains impaired in dyslexic adults, who find it difficult to recognize syllable stress patterns in words (Leong et al., 2011). Speech rhythm processing remains impaired in adults even after dyslexic individuals have developed sufficiently strong reading skills to attend university (Keshavarzi, 2025). Adult university students with dyslexia also continue to exhibit impaired sensitivity to amplitude envelope ‘rise time’ (an acoustic correlate of rhythm; Pasquini et al., 2007; Bishop-Liebler et al., 2014), a sensory impairment also found in children with dyslexia across languages (Goswami, 2015, for review). The current BCI for dyslexia was thus developed to remediate the unconscious neural factors associated with inefficient phonological development by Temporal Sampling theory.

Temporal Sampling theory accounts for the origins of the phonological ‘core deficit’ by proposing that sensory/neural processing differences regarding speech rhythm patterns cause affected individuals to develop atypical phonological representations of spoken words, which affects all levels of the linguistic hierarchy (prosodic feet, syllables, rhymes, phonemes, see Goswami, 2022b). It is known that speech is encoded by neuroelectric oscillations (rhythmic changes in electrical brain potentials in large cell networks) which respond to different temporal levels of speech information (such as phrases, syllables and phonemes, Giraud & Poeppel, 2012; Gross et al., 2013). In developmental dyslexia, Temporal Sampling theory proposes that encoding of low-frequency envelope information <10 Hz (delta and theta band information) is impaired. This arises in part because of impaired auditory discrimination of amplitude ‘rise times’ in the speech amplitude envelope. The amplitude envelope is the slow-varying energy contour of speech that determines the perception of speech rhythm (Greenberg, 2006), and it contains a range of amplitude modulation patterns at different temporal rates (Leong & Goswami, 2015). These rates broadly match EEG signalling rates such as delta, theta and beta/low gamma (∼12 – 30 Hz). ‘Rise times’ in amplitude (the rates of change between sound onset and sound peak in a given amplitude modulation) provide sensory landmarks that automatically trigger brain rhythms and speech rhythms into temporal alignment, via phase-resetting ongoing neural activity (Doelling et al., 2014). For low-frequency envelope information, neural studies suggest that this phase-resetting process is impaired in both adults and children with dyslexia, across languages (Power et al., 2016; Molinaro et al., 2016; Di Liberto et al., 2018; Destoky et al., 2020; Lizarazu et al., 2021).

Temporal Sampling theory thus proposes that an emergent phonological system can be extracted from the speech signal via the *automatic alignment* of neuroelectric oscillations to the amplitude modulation information in the speech signal, as long as amplitude rise time discrimination is effective. When amplitude rise times are not discriminated accurately, this automatic speech-brain alignment process will be negatively impacted, and the phonological system will develop less accurate representations of the sound structure of words. As phonological learning begins in infancy, infant and child EEG (electroencephalography) studies of neural speech processing informed the design of the current BCI. When infants listen to sung infant-directed speech, which is highly rhythmic, cortical tracking of low-frequency speech envelopes in the delta and theta electrophysiological bands (0.5 – 4 Hz, 4 – 8 Hz) underpin their speech processing (Attaheri et al., 2022; Ni Choisdealbha et al., 2023). Delta– and theta-band cortical tracking also underpins the learning of phonetic information by infants (Di Liberto et al., 2023). Individual differences in delta-band cortical tracking at 11 months and in the ratio of theta-delta PSD (power spectral density) predict individual differences in phonological development in early childhood (Attaheri et al., 2024). Studies of English-speaking children with developmental dyslexia also show that both the accuracy of delta-band cortical tracking and the theta-delta oscillatory ratio during natural speech listening are related to phonological development (Power et al., 2016; Di Liberto et al., 2018; Keshavarzi et al., 2022; Araújo et al., 2024). For delta-band cortical tracking, impairments are also found for dyslexic children who speak Spanish or French (Molinaro et al., 2016; Destoky et al., 2020). During continuous speech listening, English-speaking children with dyslexia show a higher theta-delta ratio than control children (not yet tested in other languages). The size of the theta/delta ratio is significantly related to their offline performance in phonological awareness tasks, with a higher ratio associated with worse performance (Araújo et al., 2024). This matches the data from English-learning infants aged 4 – 11 months. Infants with a higher theta-delta ratio during continuous speech listening went on to exhibit poorer phonological skills as young children, measured by nonword repetition (Attaheri et al., 2024). Finally, when children with dyslexia listen to speech that has been filtered to enhance the delta-band envelope information and exaggerate amplitude rise times, the theta-delta ratio normalizes (Mandke et al., 2023). This recent developmental research suggests that an individual’s theta-delta ratio during natural speech listening could be an effective target for a BCI for dyslexia which targets phonology.

The aim of the current BCI was to change the ratio of the neural oscillations that (by Temporal Sampling theory) underpin statistical learning of the amplitude modulation hierarchy in natural speech, thereby targeting the ‘phonological deficit’ and improving word reading. A non-invasive BCI focused on the self-regulation of low-frequency (delta and theta) neural oscillations during natural speech listening was thus developed (Araújo, 2023; Araújo et al., 2023). This closed-loop operant learning BCI was piloted with 15 adult participants during the Pandemic, 7 of whom had a statement of dyslexia (Araújo, 2023). The BCI was based on a spaceship flying upwards on a computer screen, and BCI players learned to self-regulate their own theta-delta ratio by controlling the spaceship’s position (Araújo, 2023). Successful BCI learning was indexed by whether the spaceship position distribution of session 1 had a significantly lower mean than session 16 (using a *t*-test, see Araújo, 2023). This *t* statistic (BCI score) reflected the degree to which the participant had reduced their theta-delta ratio. Araújo (2023) reported that 80% of participants (both dyslexic and control) learned the BCI, and that this was achieved by significantly increasing neural signal magnitude for the slower delta rhythm and significantly decreasing it for the faster theta rhythm. Further, individual BCI scores were associated with significant improvement in the speed of syllable stress discrimination judgements (phonology, *r*= 0.59, *p*< .05) and a trend in improvement for single word reading (Test of Word Reading Efficiency [TOWRE], Torgesen, Wagner & Rashotte, 1999; *r*= 0.48, *p*= .07). Adult participants also received a test of arithmetic reasoning (WRAT, Wide Range Achievement Test, Snelbaker et al., 2001) during the study, and as expected BCI scores were not associated with changes in arithmetical reasoning (Araújo, 2023). Given these promising pilot data, the BCI was adapted for mobile EEG headcaps and a further exploratory study was conducted (described here). This was necessary as Araújo’s (2023) original BCI was based on a *g-tec* hardware set-up which was not suitable for taking to schools and using with children in future studies.

There are also some BCIs in the literature which have been developed on the basis of visual and attentional theories of dyslexia (Günet, 2020; Christodoulides et al., 2022), of which one has been used with children. Günet (2020) used a commercial EEG device (Emotiv EPOC+) to create a BCI based on multisensory training of attention and vision for Turkish children with dyslexia. In her 10-minute protocol, the BCI was aimed at reducing theta activation. Children saw an arrow on a computer screen which was red, and were told to try and turn it green with brain power. If theta power was successfully reduced, the arrow would turn green. Children also received a 10-minute session of multisensory learning of the alphabet after each BCI session. Günet (2020) did not report any specific effects of the BCI protocol on theta activation in her PhD. However, the children in the BCI group made equivalent progress in reading and phonological awareness measures to control children with dyslexia who received teaching from a special education therapist. Given the experimental design, it was not possible to disentangle potential effects of any reduction in theta that may have been achieved by the BCI from the effects of the non-BCI multisensory training about letters.

Christodoulides et al. (2022) used the same commercial EEG device (Emotiv EPOC+) to record brain activity in adults during a word recognition task, in a research design informed by the magnocellular theory of dyslexia. They reported that a visual protocol with music in the background was most successful in classifying 26 Greek adult university students into two groups, a group of 14 with dyslexia and a group of 12 controls. Classification was achieved on the basis of whole brain activity across all EEG frequency bands. Their study did not aim to improve reading or phonology.

Closest to the BCI developed here, because it is also based on Temporal Sampling theory, is a BCI for dyslexia being developed by Arias et al. (2021) that is intended to improve cortical tracking of natural speech in the delta and theta EEG bands. Arias et al. (2021) describe the development of several toolboxes intended to support their BCI, which is based on computing phase locking values between the speech signal and brain activation in these EEG bands. So far, to our knowledge, this BCI has yet to be tested with participants with dyslexia.

The BCI described here was based on the BCI developed during the Pandemic by Araújo (2023). A new protocol was developed as Araújo’s original BCI was based on a *g-tec* hardware set-up which was not suitable for eventually taking to schools and using with children. Accordingly, as a further pilot, in the current study the closed-loop operant learning system was adapted to work with mobile EEG headcaps, specifically the CGX Quick-20m wireless headset. This system employs dry electrodes recorded from electrodes positioned at 19 scalp locations following the International 10-20 system (P1, FP2, F3, F4, Fz, F7, F8, C3, C4, T3, T4, T5, T6, P3, P4, Pz, O1, and O2), with A1 and A2 serving as linked-ear references. Signals were digitized at 24-bit resolution and sampled at 500 Hz. The portability and ease of setup of this dry electrode system make it particularly well suited for eventual use in schools with children, where the application of traditional EEG systems would be impractical. A new group of adults with and without dyslexia were recruited, and received a similar protocol to that used in Araújo (2023), which is described fully below. Participants were pre– and post-tested on a range of phonological, reading and control tasks (detailed below). The hypothesis was that learning the BCI would improve their neural theta-delta ratios during natural language listening, and that this improvement (indexed by their BCI scores based on spaceship position) would be significantly associated with improvements in reading and phonological processing, but not mathematics.

## 2. Materials and Methods

### 2.1. Participants

Twelve control (typically-developing) adults (mean age of 24.21 ± 6.31 years; 7 female and 5 male) and twenty adults with a current or childhood diagnosis of dyslexia (mean age of 23.65 ± 5.52 years; 16 female and 4 male) participated in the study. Nine of the 20 dyslexic participants had been diagnosed in childhood (before the age of 18 years) and the rest had received diagnoses as adults. The mean age of diagnosis in childhood was 12 years 11 months. All participants were native English speakers with normal or corrected-to-normal vision and no reported hearing impairments. Typically developing participants were included if their efficiency index (EI) on the TOWRE exceeded 95 (the EI mean is 100, S.D. 15, see Section 2.2 for further detail). Participants in the dyslexia group were included only if they could provide formal documentation of a dyslexia diagnosis from a qualified professional, such as a Health and Care Professions Council (HCPC) registered assessor, the Accessibility and Disability Resource Centre at University of Cambridge, or a specialist teacher with a current Specific Learning Difficulties (SpLD) Assessment Practicing Certificate. This preceding documentation was scrutinized to ascertain possible co-morbid diagnoses such as ADHD, as well as the nature of the child’s cognitive difficulties that had led to a formal diagnosis. None of the 20 dyslexic participants had a formal diagnosis of ADHD, and there was no prior documentation of possible visual or magnocellular difficulties. Accordingly, all participants were assumed to have developmental phonological dyslexia. We also asked our participants about co-morbid disorders prior to participating in the BCI study. One participant mentioned that she had self-diagnosed ADHD. All participants provided informed consent for the study in accordance with the Declaration of Helsinki, and the study was reviewed by the Psychology Research Ethics Committee of the University of Cambridge who gave it a favourable opinion (PRE.2022.034).

### 2.2. Experimental Protocol

The experimental protocol spanned ten days and included two assessment sessions, one at the beginning (Day 1) and one at the end (Day 10), to evaluate participants’ cognitive and linguistic profiles. These pre– and post-intervention sessions comprised tasks measuring phonological and reading skills, acoustic processing, and skills that were not expected to be improved by the BCI (non-verbal reasoning and arithmetic ability). The eight days in between were dedicated to the BCI training intervention, which will be described in Section 2.3. The experimental protocol can be found in Figure 2.1(a). The measures used in the pre– and post-test sessions are described below.

**Figure 2.1.**
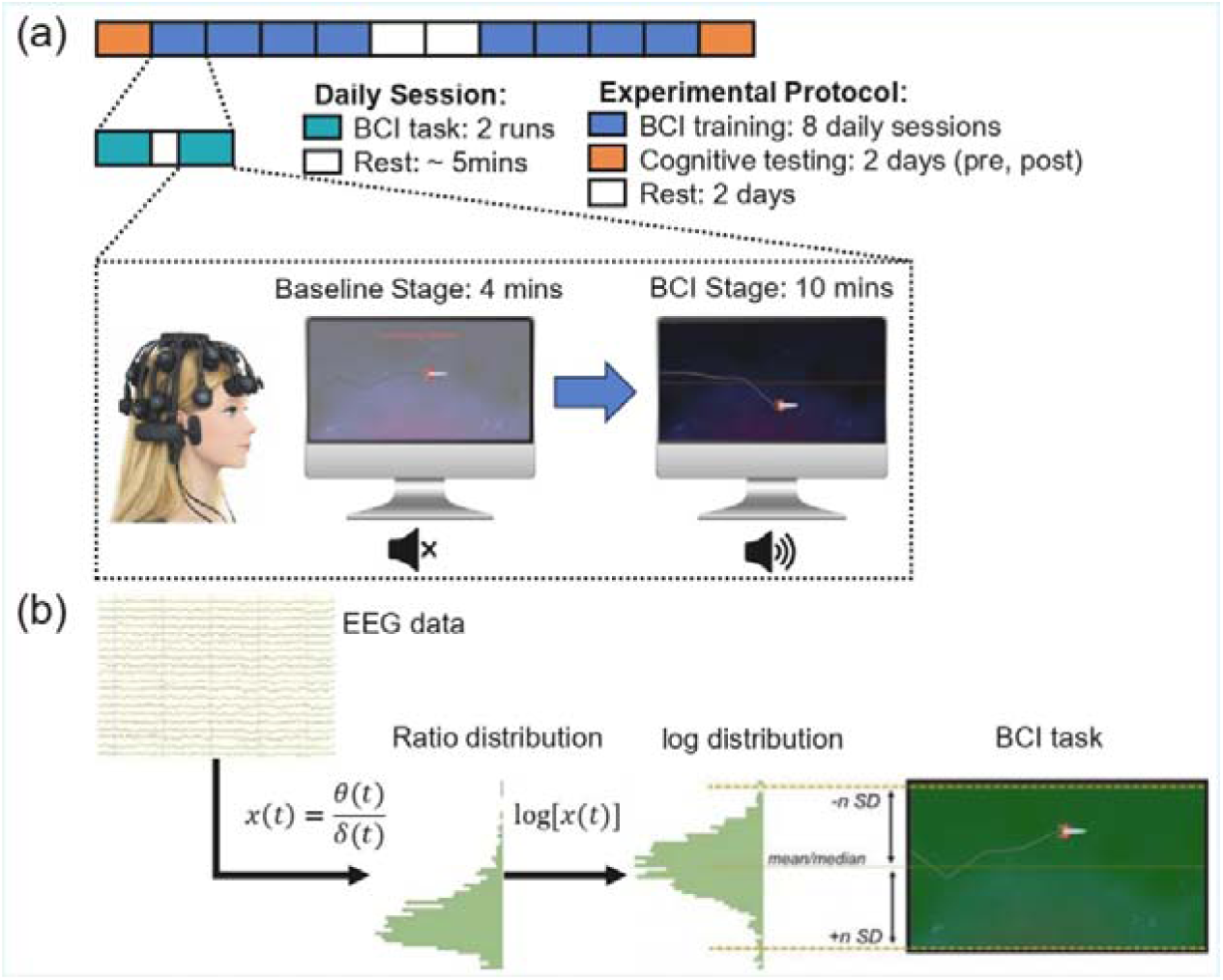
Panel (A) shows the whole Experimental Protocol, including the time line of each BCI session and the spaceship interface. Panel (B) depicts the decoder logic behind the neural feedback.

#### 1) Phonology and Reading Measures

Phonology and reading skills were measured using an experimental phoneme deletion task, an experimental syllable stress recognition task, an experimental Rapid Automatized Naming (RAN), and the standardized word and nonword item lists from the TOWRE.

The phoneme deletion task was adapted from McDougall et al. (1994). This task required participants to listen to a spoken item and delete a target phoneme (e.g., “BICE” without the /b/ becomes “ICE”). The target consonant phoneme appeared in initial, medial, or final positions, and all correct responses formed real English words. The task comprised 18 trials (3 practice and 15 experimental items), presented through sound files recorded by a female speaker of standard Southern British English. The same test was administered both before and after the BCI training.

In the syllable stress discrimination task, participants heard pairs of different four-syllable words and made a same-different judgement regarding whether the pair of words shared the same stress pattern (e.g., difficulty – voluntary = yes). The items were from Leong et al. (2011), and comprised 80 randomized pairs, some of which were deliberately mis-stressed (e.g., di-FFI-cul-ty – VO-lun-ta-ry = NO). This task was administered before and after the BCI training. This task was a variation to the protocol used in Araújo (2023), in which pairs of identical words were used as stimuli (also drawn from Leong et al., 2011). Here we used comparisons between different words to increase task difficulty, with the aim of reducing the ceiling effect observed with original design.

In the RAN task, participants named pictures of familiar items (e.g., cup, book, tree) aloud as fast as possible. Four pages of pictures were administered at both the pre– and post-sessions. Two pages of target pictures represented words with low phonological neighborhoods (such as ‘cup’ and ‘book’, which have few similar-sounding items in the mental lexicon) and two pages of target pictures represented words with high phonological neighborhoods (such as ‘wheel’ and ‘gate’, which have many similar-sounding items in the mental lexicon, see De Cara & Goswami, 2002). Both the time taken to complete the task and accuracy were recorded, and an averaged score for response time and accuracy for the 4 pages of items was utilised in the analyses.

Reading was assessed using the TOWRE. The participant received a list of single words to read aloud in 45 seconds, and a list of nonword items to read aloud in 45 seconds. Version A of this task was given at the pre-training stage while version B was used at the post-training stage. The highest available age bracket for calculating scaled scores ranges from 17 years 0 months to 24 years 11 months. Since some participants in the study were older than this range, raw scores were used for statistical analyses. However, for participant recruitment, an EI was calculated using scaled scores from 17-24 age group, as all participants were over 18 years of age. These scaled scores were used for typically developing group screening purposes and were not included in further analysis

#### 2) Non-verbal I.Q

All participants completed the matrix reasoning subtest of the Wechsler Intelligence Scale for Adults (WAIS; Wechsler, 1955), a widely used measure of non-verbal intelligence. This is a nonverbal reasoning task in which individuals are asked to identify patterns in designs. This pattern recognition task was administered at both pre– and post-intervention sessions.

#### 3) Arithmetic Task

Participants completed the standardized arithmetic subscale of the Wide Range Achievement Test (WRAT) (Snelbaker et al., 2001), which includes basic math problems requiring written responses. Version TAN was administered before the intervention, and Version BLUE after. This task was included to test whether the BCI would affect any academic skill, rather than specifically affect word reading.

#### 4) Acoustic Threshold for Amplitude Rise Time (ART): 1 Rise Task

Participants also completed a sine tone rise time task (labelled the 1 Rise task in our prior publications with children, e.g. Flanagan et al., 2024) to assess sensitivity to ART. Each trial presented three 500-Hz tones, with one (the target) having a slower onset rise time than the two standard tones. Using an AXB format displayed as cartoon dinosaurs, participants were asked to identify which of the first or third sounds differed from the middle tone. The task used 39 stimuli with rise times ranging from 300 ms to 15 ms in 7.3 ms steps. Verbal instructions and five practice trials with feedback were provided before the main task. Participants performed the same test both before and after the BCI training.

### 2.3. BCI Training

Between the pre– and post-intervention assessment sessions, participants completed eight days of BCI training. The protocol was structured to span ten days in total, allowing at most two rest days (typically the weekend) to accommodate participant schedules. Most participants completed the intervention over two consecutive weeks. Each daily session included two BCI runs. The experimenter running the BCI was not blind to group membership. Prior to each session, an EEG cap was fitted, and electrode impedances were checked and maintained below 100 Ω. Participants were encouraged to listen carefully to the words in the story and try to identify listening strategies to keep the spaceship ascending on the screen. However, no explicit suggestions regarding how to achieve this goal were given.

Each BCI run consisted of two distinct phases: a baseline stage and a BCI control stage. The interface of the BCI and the timeline of the experiment are depicted in Figure 2.1(a). In the baseline stage (lasting four minutes), participants viewed a vertically moving spaceship displayed on the screen. During this phase, they had no neural control over the spaceship’s position. Instead, the spaceship moved randomly, with positions sampled from a Gaussian distribution (mean = 0.5, SD = 0.15) and mapped onto a vertical scale ranging from 0 (top) to 1 (bottom). The position updated at a refresh rate of 4 Hz. This random movement served two purposes: it provided data to estimate individualized decoder thresholds based on each participant’s typical neural activity, and it avoided any neural entrainment that might occur with fixed or repetitive visual patterns. A semi-transparent white overlay and the message “Good luck! Please wait…” were displayed to indicate the system was in passive mode. No auditory input was presented during this stage.

After the baseline, participants entered the BCI stage, which lasted for the duration of a ten-minute auditory story. At this point, the semi-transparent overlay and the baseline message were removed, and participants began listening to a narrated version of Winnie-the-Pooh through headphones. Winnie-the-Pooh is a simple children’s story which was easy to comprehend and did not require verbal retention of difficult material. It begins as follows “Once upon a time, a very long time ago now, about last Friday, Winnie-the-Pooh lived in a forest all by himself under the name of Sanders. “What does ‘under the name’ mean?” you might ask. It means he had the name over the door in gold letters, and lived under it….”. The same chapter of the story was used in all sessions. Simultaneously, participants gained neural control over the on-screen spaceship, which moved vertically based on real-time EEG activity. Specifically, the spaceship’s position was determined by the log-transformed theta/delta power ratio measured from centrally located electrodes (F3, F4, C3, Cz, C4, P3, P4). To personalize control sensitivity, decoder boundaries were set using each participant’s baseline distribution: the median of their log-transformed theta/delta ratio defined the vertical midline of the screen, while the upper and lower boundaries were set at three standard deviations above and below the median. Participants were instructed to raise the spaceship as high as possible and to keep it stable during the story. In terms of neural dynamics, this corresponded to decreasing the theta/delta ratio and reducing its variance.

The BCI was designed to provide feedback based on neural patterns previously associated with continuous speech processing and phonological awareness in children with and without dyslexia. The same decoder was used across participants, with calibration derived from each session’s baseline. This allowed for continuous control based on dyslexia-relevant neural dynamics, specifically those shown to relate to phonological awareness in previous studies.

The neurofeedback display was designed to be intuitive and engaging. To enhance motivation and user experience, the traditional cursor was replaced with a spaceship graphic, and the background featured a subtle space-themed design. A visual midline was drawn on the screen to indicate the target region for upward control. In addition, a five-timestep history trace was implemented, appearing as a contrail behind the spaceship, allowing participants to visually track their recent performance. To enhance participant motivation, a cumulative score related to the real-time theta/delta ratio was presented in the top-left corner of the screen. The score was updated at each refresh of the spaceship position. The instantaneous score was derived from the log transform of the ratio standardized to each participant’s baseline, which also determined the spaceship’s position. It was multiplied by 10 when the spaceship occupied the lower half of the screen and by 20 when it occupied the upper half to reinforce positive feedback. Each instantaneous score was continually added to the total score displayed. The story audio was not influenced by task performance and remained constant throughout the session.

To further reinforce successful BCI control, a visual reward system was implemented. A semi-transparent green overlay appeared on the screen, with its intensity varying according to the spaceship’s vertical position. The screen was scaled from 0 (top) to 1 (bottom), and the green glow was calculated using the formula:

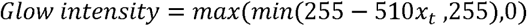

Where *x*_1_ is the scaled spaceship position. This meant that the glow reached full intensity when the spaceship was at the top of the screen, gradually faded when it approached the midline, and disappeared entirely when below the midline. This continuous visual reinforcement served as an intuitive feedback signal to encourage better control of the spaceship.

To prepare the signal for use in the BCI decoder, the theta/delta ratio was log-transformed. This transformation was necessary because the raw ratio data exhibited a skewed, non-Gaussian distribution across and within participants, along with a wide and variable dynamic range. Applying a log transformation reduced skewness and the influence of outliers by compressing the range of values. Crucially, because the log function is monotonically increasing, it preserved the relative ordering of values in the original signal. This ensured that the neurofeedback interface remained stable, interpretable, and sensitive to individual neural dynamics. The BCI neural feedback is shown in Figure 2.1(b).

### 2.4. EEG preprocessing

The EEG signal was acquired in real time using a CGX wireless headset and continuously processed throughout the neurofeedback task. Raw data were initially converted from a 24-bit compressed format to microvolts and streamed at a sampling rate of 500 Hz. EEG preprocessing followed two distinct strategies: *real-time processing* for neurofeedback delivery, and *offline preprocessing* for subsequent data analysis. The rationale for using two strategies was as follows. Real-time processing is typically the learning measure reported for BCI studies, as was the case for Araújo (2023). However with the mobile EEG headcap, online artifact rejection could not completely rule out the potential confounding effects of factors such as abnormal eye movements or subvocal rehearsal. Accordingly, an offline processing strategy was also applied, using more computationally intensive and manually supervised techniques, including Independent Component Analysis (ICA) and Artifact Subspace Reconstruction (ASR), to suppress physiological and non-physiological artifacts such as muscle bursts, eye movements, and poor electrode-skin contact. Each strategy is described in more detail below.

#### 2.4.1 Real-Time Processing Protocol

During the BCI intervention, real-time processing prioritized low computational demand and effective noise suppression to ensure smooth feedback. A zero-phase, fourth-order Butterworth bandpass filter (0.5–10 Hz) was applied to selected central channels to reduce noise and isolate relevant neural signals. A 3-second sliding window was used for continuous feature extraction. During pilot testing, we observed that the spaceship position could be influenced by abnormal eye movements. To manage transient artifacts (e.g., eye movement, muscle activity or movement), an online threshold-based artifact rejection method was employed. Samples exceeding a channel-specific threshold—determined from the 95th percentile of the participant’s baseline amplitude distribution—were replaced with random clean segments drawn from the individual’s baseline data, preserving inter-channel relationships. This procedure ensured that the spaceship position was less affected by abnormal EEG segments during BCI training. The resulting preprocessed signal was then used for real-time spectral analysis and neurofeedback computation.

#### 2.4.2. Offline Preprocessing Protocol

Despite the use of an online artifact rejection method, the influence of artifacts on spaceship position could not be completely eliminated during online processing. To address this limitation, a more comprehensive offline preprocessing pipeline was applied to obtain cleaner data. Power line noise was removed using a notch filter, and the data were bandpass-filtered between 0.5 and 48 Hz using an 8th-order Butterworth filter with zero-phase filtering to avoid phase distortion. The signal was then downsampled to 250 Hz to reduce computational load. Given that dry EEG systems tend to produce noisier recordings than gel-based systems, ASR was used to suppress high-amplitude transient artifacts such as muscle bursts (e.g., from eye movements or subvocalization). Channels were marked as noisy if their voltages exceeded ±100 μV or if their power spectra deviated more than 3 standard deviations from the mean. These noisy channels were excluded from the computation of the average reference and from ICA fitting. On average, 0.62 ± 0.71 channels were marked as noisy at this stage. The EEG data were then re-referenced to the average of the remaining non-noisy channels.

ICA was then performed, and components associated with ocular, muscular, or blink artifacts (e.g., EOG, EMG) were identified and removed manually. On average, 3.43 ± 1.70 components were removed per recording. The cleaned data were segmented into consecutive, non-overlapping 3-second epochs. Finally, epoch-level noise rejection and interpolation were applied. A 3-second epoch was rejected if 5 or more of the 19 channels exceed ±80 µV. If 1 to 4 channels exceed ±80 µV within an epoch, the epoch is retained, and the contaminated channels were interpolated using a spline interpolation method. Following epoching, 0.52% ± 2.37% of trials were removed, and 2.39% ± 3.24% of channel-by-epoch items were interpolated.

### 2.5. Statistical Analysis

Statistical analyses were conducted to evaluate the effectiveness of the BCI neurofeedback intervention and its relationship with behavioral performance. First, to assess whether participants exhibited neurophysiological changes across the training period, we tested whether there was a significant change in participants’ neural responses over time. Specifically, we compared the distribution of the log-transformed theta/delta ratio between the first session and the final session. This was done separately for both (i) the real-time ratio used for intervention feedback (derived from the online preprocessing pipeline), and (ii) the ratio extracted from the offline preprocessed data. As noted, the offline theta/delta ratio can be considered the purer measure of BCI learning in terms of artifact removal. T-tests were used to assess differences in the ratio values across sessions for each participant. The resulting t-statistic served as a summary measure of change in BCI performance over time. Participants who showed a significantly lower ratio by the final session were considered to have demonstrated learning. To strengthen the assessment of BCI learning, Bayesian analyses were additionally performed. Bayes factors were calculated to quantify the strength of evidence for a session-related change, and Bayesian credible intervals were estimated for the mean difference between the beginning and final sessions. The posterior probability of improvement was also calculated as the probability that the beginning-session ratio exceeded the final-session ratio.

Second, to evaluate whether participants improved on relevant behavioral skills following the BCI intervention, pre– and post-intervention behavioral scores (e.g. phonological awareness, reading ability) were compared using paired-sample t-tests. A significant increase in post-test scores was interpreted as evidence of behavioral improvement. Pre– and post-test performance was computed both for the entire sample (as some learning of the BCI may have occurred even in participants whose BCI *t-*statistic was not significant) and separately for participants who demonstrated BCI learning according to each protocol (real-time versus offline artifact removal).

Third, we investigated whether changes in neural measures were associated with changes in behavioral performance. Given the small sample and possible sensitivity to outliers, bend correlations were conducted to assess the relationship between the t-statistics derived from the BCI measures (both real-time and offline) and the differences in behavioral performance between pre-test and post-test. The stability of each association was further evaluated using 95% bootstrap confidence intervals. For all analyses described above, p-values were corrected for multiple comparisons using the false discovery rate (FDR), and statistical significance was defined as corrected p < 0.05.

## 3. Results

### 3.1. Neurophysiological Changes Across Sessions

To evaluate whether participants exhibited neurophysiological changes across the BCI training period, we compared the log-transformed theta/delta power ratios between the first and final sessions. This analysis was conducted twice, separately for data processed using the real-time processing pipeline (following Araújo, 2023) and for data pre-processed offline (more stringent analysis added here).

For each participant, a *t*-test was conducted to assess whether the theta/delta ratio significantly decreased from the first to the last session. The resulting *t*-statistic served as an individual-level summary of neural change and was used as the participant’s *BCI learning score* (hereafter BCI score) in subsequent analyses. Participants were considered to have learned the BCI control task successfully if their *t*-statistic was greater than zero and the corresponding *p*-value was less than 0.05. This criterion indicates a statistically significant reduction in the theta/delta ratio across BCI sessions.

Based on the **real-time** EEG data (artifacts removed in real time), 9 out of 12 participants in the control group and 13 out of 20 participants in the dyslexia group met this learning criterion. Following offline preprocessing, which involved the removal of ocular, muscular, and movement-related artifacts, the resulting theta/delta ratios exhibited generally lower *t*-statistics. Under this more stringent **offline** preprocessing protocol, 6 of 12 participants in the control group and 10 of 20 in the dyslexia group showed significant improvement according to the same criterion. Accordingly, half of the participants in each group showed learning of the BCI following stringent artifact removal.

To further quantify the strength and direction of evidence for learning, directional Bayes factors were calculated for the reduction in offline theta/delta ratio from the first to the last session. A *BF*_10_ greater than 1 indicated evidence in favour of the hypothesised reduction over the null hypothesis, whereas a *BF*_10_ below 1 indicated evidence in favour of the null hypothesis. Following Lee and Wagenmakers (2014), values between 1 and 3 were interpreted as anecdotal evidence, 3–10 as moderate evidence, and >10 as strong evidence for the hypothesised reduction. Bayesian results were broadly consistent with the t-test classification. In the control group, all six participants classified as learners showed strong evidence for improvement, with *BF*_10_ values greater than 1 ranging from 41.14 to 5.69 × 10^48^, and 95% credible intervals for the first-minus-last session difference excluding zero in the positive direction. The remaining six control participants showed *BF*_10_ values below 1, indicating the data favoured the null hypothesis; five of these participants had credible intervals excluding zero in the opposite direction, while one had a credible interval overlapping zero. In the dyslexia group, nine participants classified as learners showed *BF*_10_ values greater than 3 and credible intervals excluding zero in the expected positive direction. Of these learners, one dyslexic participant showed anecdotal evidence for improvement (*BF*_10_ = 1.27), while the other nine showed strong evidence for improvement, with *BF*_10_ ranging from 9.75 × 10^6^ to 2.75 × 10^46^. Among the ten dyslexia participants classified as non-learners, all showed *BF*_10_ values below 1. Seven showed credible intervals excluding zero in the opposite direction, while three showed inconclusive evidence, with credible intervals overlapping zero. Figure 3.1(A) presents the log_10_ *BF*_10_ distribution of offline theta/delta ratio from the both groups. *BF*_10_ values were log-transformed for ease of interpretation.

**Figure 3.1.**
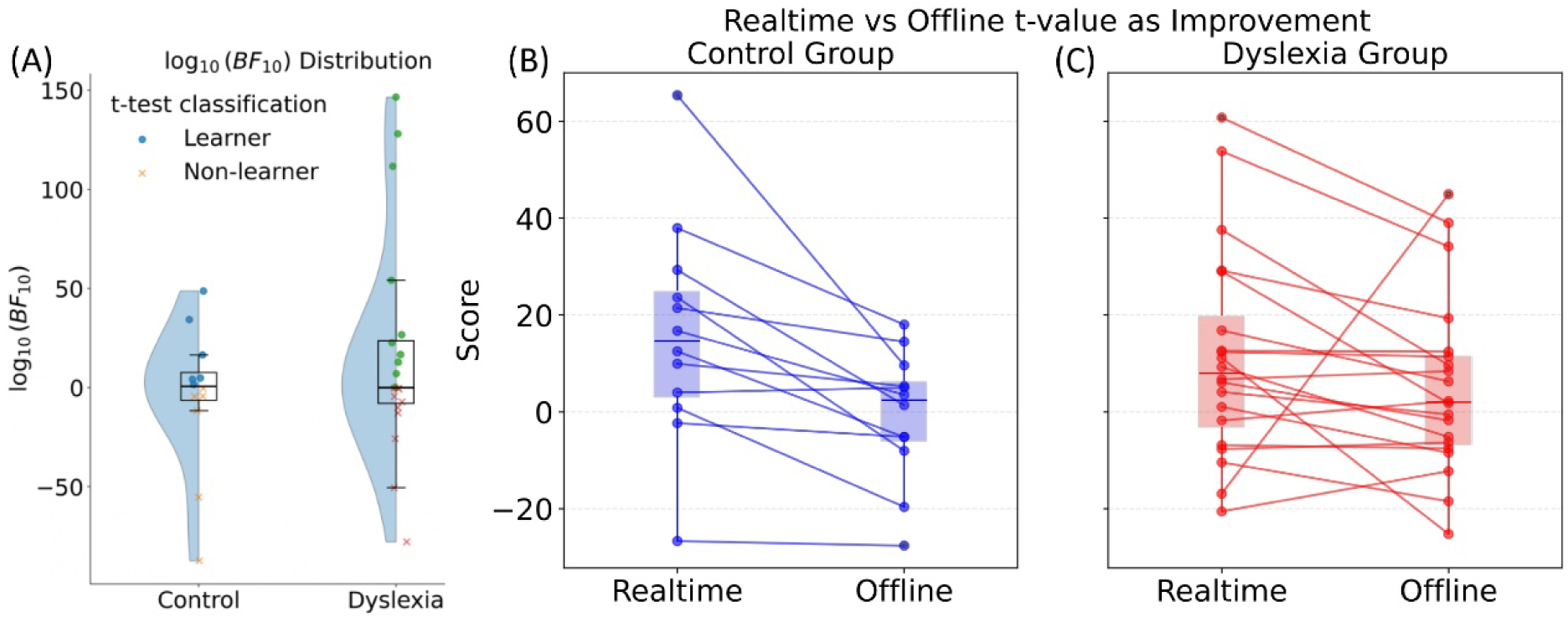
(A) Distribution of participant-level log_10_ *BF*_10_ values for the control and dyslexia groups. Half-violin plots show the estimated distributions, boxplots summarise the median and interquartile range, and individual points represent participants. (B-C) The *t*-statistics used as a measure of BCI performance. Data are shown for each participant, under real-time feedback versus offline preprocessing. The left subplot (3B, in blue) represents the control group, and the right subplot (3C, in red) represents the dyslexia group. In each subplot, individual participant data are shown using paired line plots that connect the t-statistics obtained from the real-time and offline pipelines, illustrating the direction and magnitude of change.

Figure 3.1(B) and (C) presents a visual summary of the *t-*statistics for each participant under both processing pipelines. As the offline preprocessing pipeline is more stringent, it would be expected that the BCI performance scores are lower for the offline preprocessing, which was the case for both groups. Overall, a higher *T*-score indicates better learning of the BCI. Overlaid on each set of lines, boxplots depict the overall distribution of *t*-statistics for each preprocessing method within each group.

### 3.2. Behavioral Improvements Following BCI Intervention

To investigate whether participants showed behavioral improvement over the course of training, we examined performance across the acoustic, cognitive and linguistic tasks. As will be recalled, two measures were not expected *a priori* to show improvement following BCI training, nonverbal IQ (WAIS Matrices) and Arithmetic.

#### 3.2.1. Whole Sample: Comparison of Pre-Intervention Behavioural Scores by Group

As a first step, we compared pre-intervention scores between the control and dyslexia groups to assess baseline differences in task performance (Table 3.1). As expected, there were no significant group differences in non-verbal cognitive tasks such as Arithmetic and Matrices Reasoning, indicating that both groups were matched on non-reading academic performance and general reasoning ability. In contrast, significant group differences were observed in pre-test measures of reading and phonological processing. These included syllable stress discrimination accuracy, RAN completion time, and both word and non-word raw scores on the TOWRE. These results are consistent with the known language and literacy difficulties associated with dyslexia. An exception was the phoneme deletion task, which showed no significant group difference. This was likely due to a ceiling effect. The task may have been too easy for both groups, limiting its sensitivity to detect individual differences. Contrary to our prior adult studies, no significant group difference in sensitivity to ART was observed, although the dyslexic group showed worse performance.

**Table 3.1.**
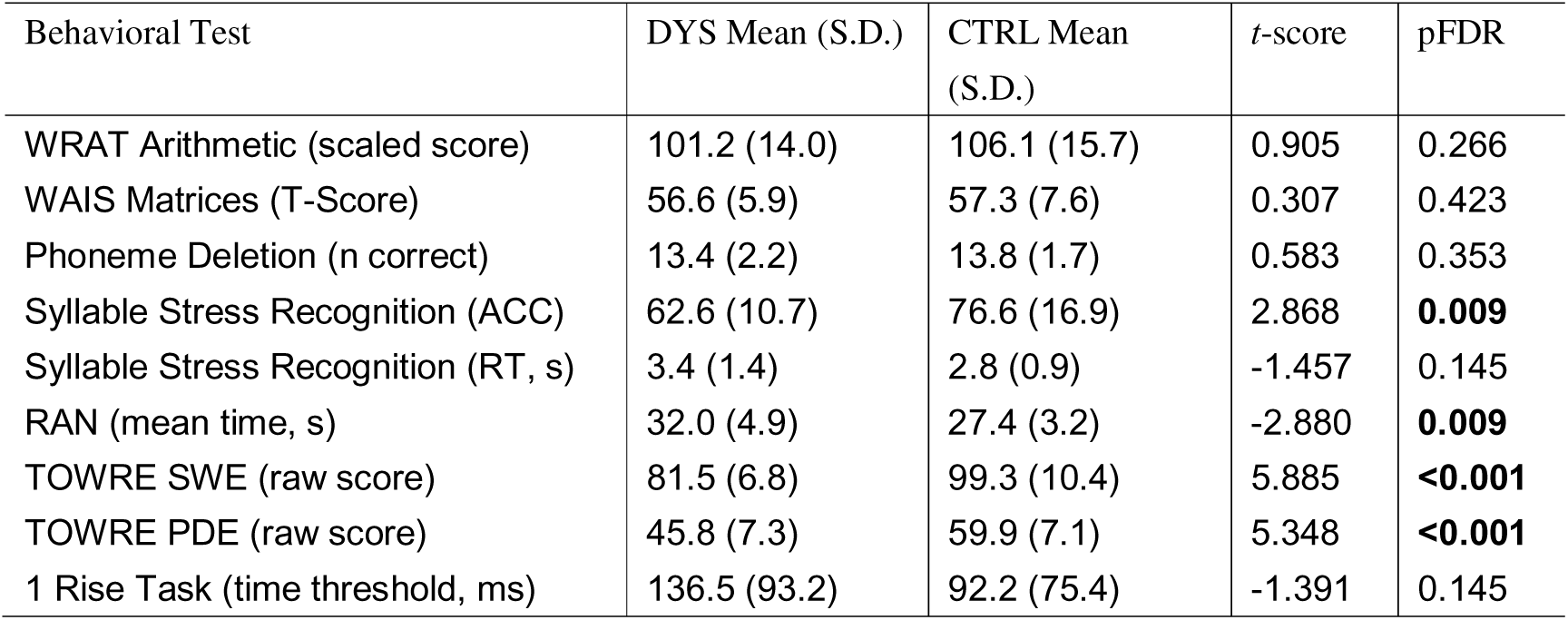
Performance on pre-test behavioural measures for the whole sample, separated by group, 2-tailed *t*-tests.

| Behavioral Test | DYS Mean (S.D.) | CTRL Mean (S.D.) | *t*-score | pFDR |
| --- | --- | --- | --- | --- |
| WRAT Arithmetic (scaled score) | 101.2 (14.0) | 106.1 (15.7) | 0.905 | 0.266 |
| WAIS Matrices (T-Score) | 56.6 (5.9) | 57.3 (7.6) | 0.307 | 0.423 |
| Phoneme Deletion (n correct) | 13.4 (2.2) | 13.8 (1.7) | 0.583 | 0.353 |
| Syllable Stress Recognition (ACC) | 62.6 (10.7) | 76.6 (16.9) | 2.868 | **0.009** |
| Syllable Stress Recognition (RT, s) | 3.4 (1.4) | 2.8 (0.9) | -1.457 | 0.145 |
| RAN (mean time, s) | 32.0 (4.9) | 27.4 (3.2) | -2.880 | **0.009** |
| TOWRE SWE (raw score) | 81.5 (6.8) | 99.3 (10.4) | 5.885 | **<0.001** |
| TOWRE PDE (raw score) | 45.8 (7.3) | 59.9 (7.1) | 5.348 | **<0.001** |
| 1 Rise Task (time threshold, ms) | 136.5 (93.2) | 92.2 (75.4) | -1.391 | 0.145 |

#### 3.2.2. Whole Sample: Comparison of Pre-Intervention versus Post-BCI Behavioural Scores by Group

We next compared pre– and post-intervention scores within each group to examine behavioral changes over time (Table 3.2). The whole sample was used because they had all experienced the BCI, even though some participants did not show statistical evidence of significant learning. In the control group, a significant improvement was observed in RAN completion time, while Arithmetic, syllable stress accuracy, and TOWRE non-word reading showed numerical improvement but did not reach significance. In the dyslexia group, significant improvements were found in Matrices Reasoning, phoneme deletion accuracy, syllable stress discrimination accuracy, RAN completion time, and both TOWRE word and non-word reading. With the exception of the improvement in Matrices Reasoning (dyslexics only), these improvements were in line with our *a priori* expectations. However, as noted both groups in these analyses included participants who did not meet criterion for learning the BCI.

**Table 3.2.** Pre– versus post-intervention scores for the behavioral tasks for the whole sample, separated by group, 2-tailed *t*-tests.

| Behavioral Test | Group | Pre-test<br>mean (S.D.) | Post-test<br>mean (S.D.) | *t*-score | *pFDR* |
| --- | --- | --- | --- | --- | --- |
| WRAT Arithmetic (scaled score) | CTRL | 106.1 (15.7) | 110.4(17.0) | -2.738 | 0.051 |
|  | DYS | 101.2 (14.0) | 102.8(14.6) | -1.281 | 0.108 |
| WAIS Matrices (T-Score) | CTRL | 57.3 (7.6) | 59.2 (7.1) | -1.675 | 0.164 |
|  | DYS | 56.6 (5.9) | 59.5 (4.9) | **-4.681** | **<0.001** |
| Phoneme Deletion (n correct) | CTRL | 13.8 (1.7) | 14.0 (1.3) | -0.561 | 0.521 |
|  | DYS | 13.4 (2.2) | 13.9 (2.1) | -2.127 | **0.031** |
| Syllable Stress Recognition (ACC) | CTRL | 76.6 (16.9) | 80.7 (18.2) | -2.865 | 0.051 |
|  | DYS | 62.6 (10.7) | 67.4 (12.6) | **-2.623** | **0.013** |
| Syllable Stress Recognition (RT, s) | CTRL | 2.8 (0.9) | 2.4 (0.9) | 1.670 | 0.164 |
|  | DYS | 3.4 (1.4) | 3.4 (1.9) | 0.062 | 0.423 |
| RAN (mean time, s) | CTRL | 27.4 (3.2) | 25.3 (3.2) | **4.891** | **0.004** |
|  | DYS | 32.0 (4.9) | 28.0 (4.3) | **6.631** | **<0.001** |
| TOWRE SWE (raw score) | CTRL | 99.3 (10.4) | 98.6 (8.9) | 0.581 | 0.521 |
|  | DYS | 81.5 (6.8) | 87.2 (7.3) | **-4.467** | **<0.001** |
| TOWRE PDE (raw score) | CTRL | 59.9 (7.1) | 61.7 (5.0) | -2.399 | 0.071 |
|  | DYS | 45.8 (7.3) | 48.4 (7.4) | **-2.931** | **0.009** |
| 1 Rise Task (time threshold, ms) | CTRL | 92.2 (75.4) | 75.2 (57.2) | 1.073 | 0.350 |
|  | DYS | 136.5 (93.2) | 119.3 (93.9) | 1.440 | 0.095 |

#### 3.2.3 Pre-Intervention versus Post-BCI Behavioural Scores, BCI Learners only

We next assessed the same relationships for the BCI learners only, analysing the data for either real-time preprocessing (while playing the BCI) or offline pre-processing (potential artifacts removed offline, see Table 3.3). In the control group, significant improvements were observed in RAN completion time, for both real-time and offline preprocessing. In the dyslexia group, significant improvements were found in Matrices reasoning, RAN completion time, syllable stress discrimination accuracy and TOWRE word and non-word reading, for both real-time and offline preprocessing. With the exception of the improvement in Matrices reasoning (dyslexics only), these improvements were in line with our *a priori* expectations based on Temporal Sampling theory. In order to assess which improvements were associated with learning the BCI, we next explored the relations between individual t-statistics and behavioural improvements.

**Table 3.3.**
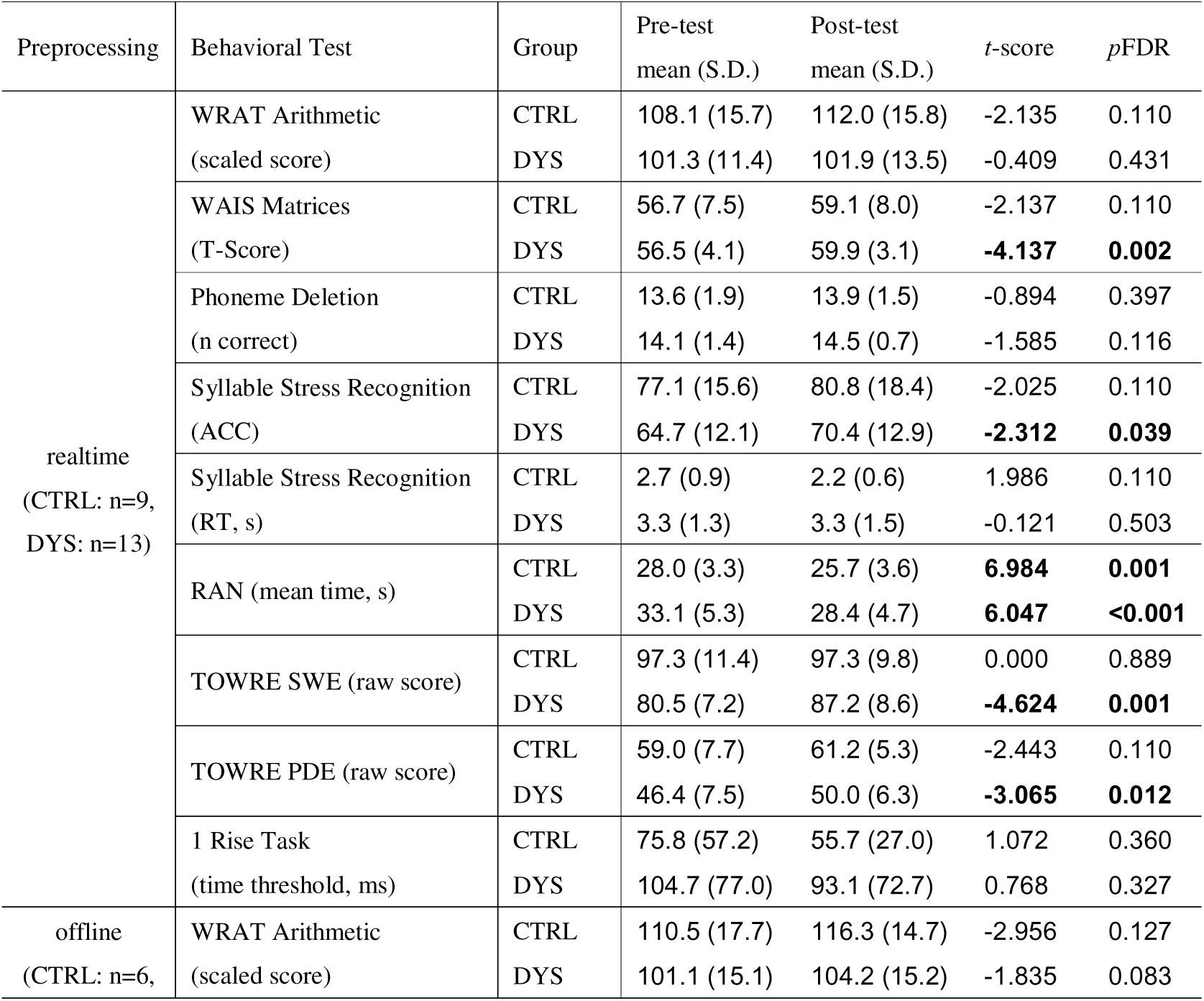
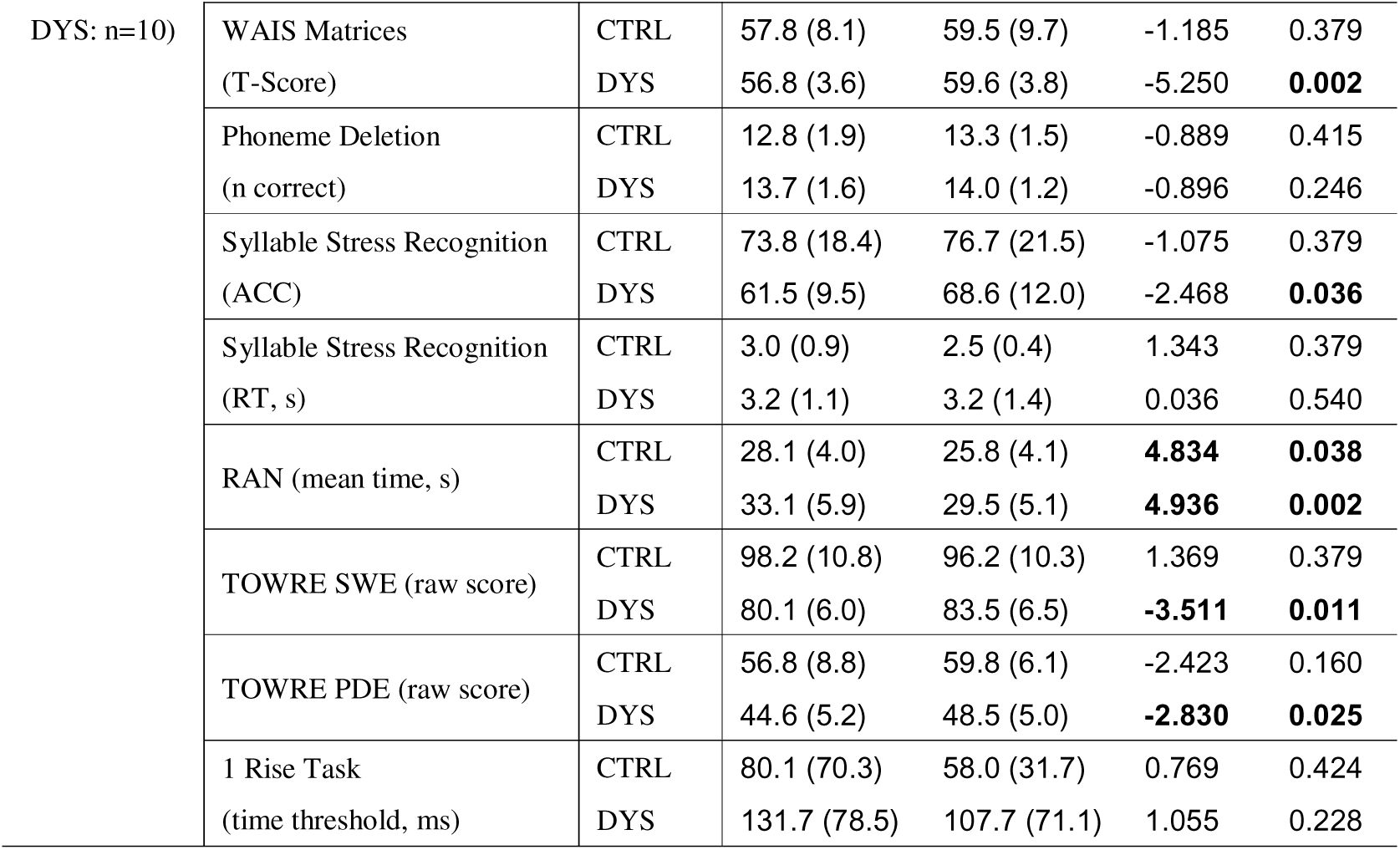
Pre– and post-intervention behavioral scores for BCI learners only by group, separated by real-time artifact removal versus more stringent offline preprocessing to remove potential artifacts; 2-tailed *t*-tests.

### 3.3. Associations Between Neural and Behavioral Changes

To explore whether the observed improvements in performance were systematically related to learning the BCI, we calculated correlations between participants’ BCI learning scores for each assessment (real-time learning, offline preprocessing) and their changes in the behavioral tasks (computed as post-intervention behavioural score minus pre-intervention score in each case). Higher *t*-statistics indicate greater success in reducing theta/delta ratios across training, and hence should be positively related to improvements in accuracy in the behavioural tasks, and negatively related to improvements in processing time. Given the relatively small sample sizes, we computed bend correlations using a bending proportion of β = 0.20 (Wilcox, 1994) instead of Pearson correlations, as the latter can be overly affected by outliers and extreme values. Given that Temporal Sampling theory would predict improvements in the phonological and reading tasks, we tested the correlations using one-tailed tests.

#### 3.3.1 All Participants: Bend Correlations with Improvements in Behavioural Tasks

The whole sample was analysed first, because all participants experienced the BCI, even though some participants did not show statistical evidence of significant learning. When the bend correlations were computed for the whole sample, only the offline EEG data (BCI learning score computed after removal of a greater number of artifacts) showed significant correlations with behavioural improvements, and for dyslexic participants only. The results are shown in Table 3.4. As can be seen, the participants with dyslexia showed significant relations between BCI learning and both phonology and reading tasks. Positive correlations were found between BCI learning scores and improvements in syllable stress discrimination accuracy (*r*= .65, FDR-corrected *p*< .005, 95% CI [.18,.85], Figure 3.2(B)), and also between BCI learning scores and TOWRE non-word reading (*r*= .51, FDR-corrected *p*< .03, CI [.06, .81], Figure 3.2(A)). In both cases, the 95% confidence intervals were entirely positive and did not include zero, supporting the reliability of these positive associations. This suggests that when BCI learning was assessed with the more stringent preprocessing protocol, the BCI had the intended effect of improving phonological processing and improving phonological recoding of print to sound in participants with dyslexia. The relevant scatter plots and regression lines are shown in Figure 3.2 (offline preprocessing).

**Figure 3.2.**
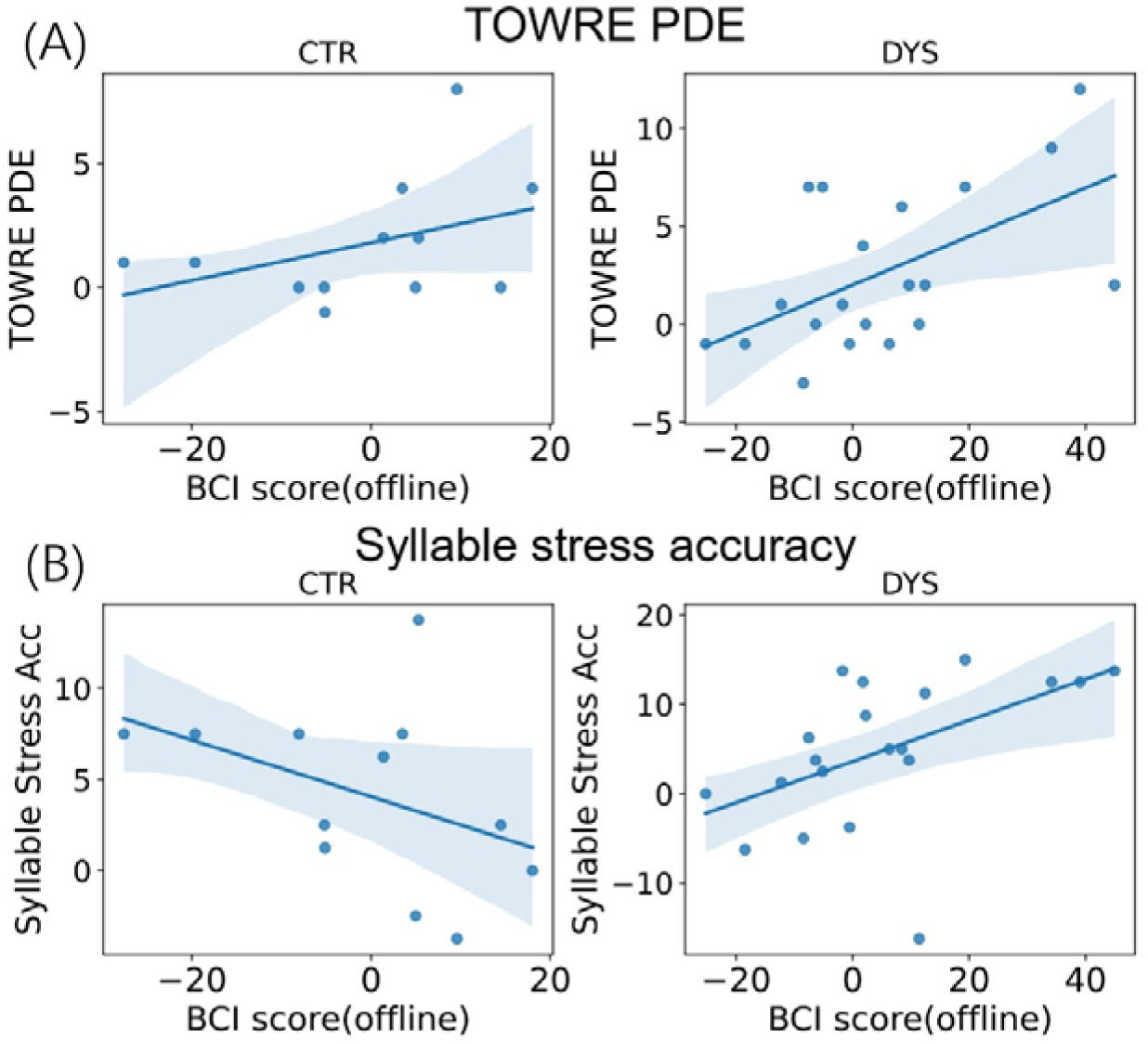
Scatterplots depicting relation between BCI scores from offline preprocessing and behavioural task scores in the full sample, shown separately for control and dyslexia groups. Behavioural measures are (A) TOWRE nonword reading and (B) syllable stress perception accuracy. Lines show the linear fit for visualisation.

**Table 3.4.**
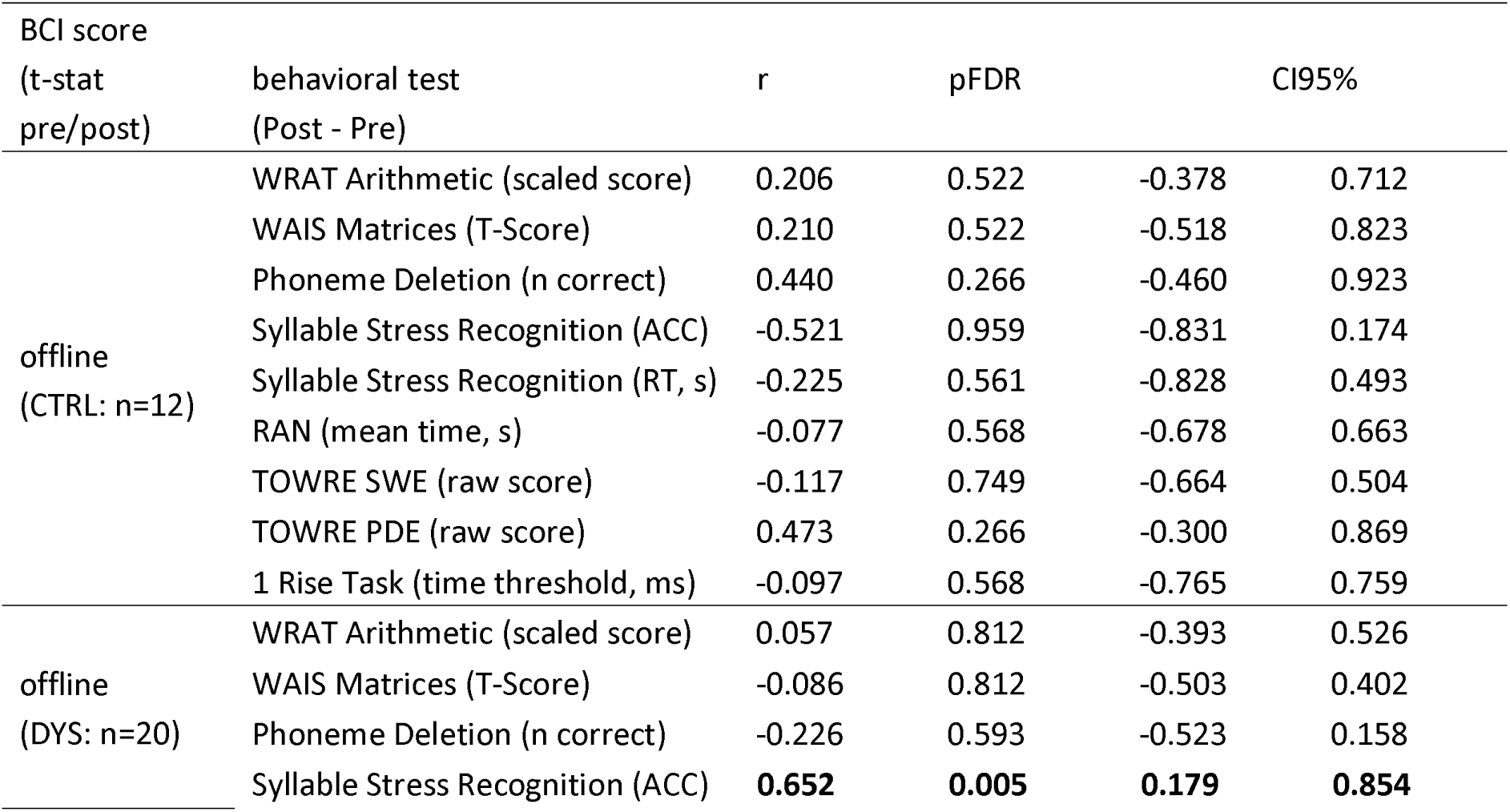
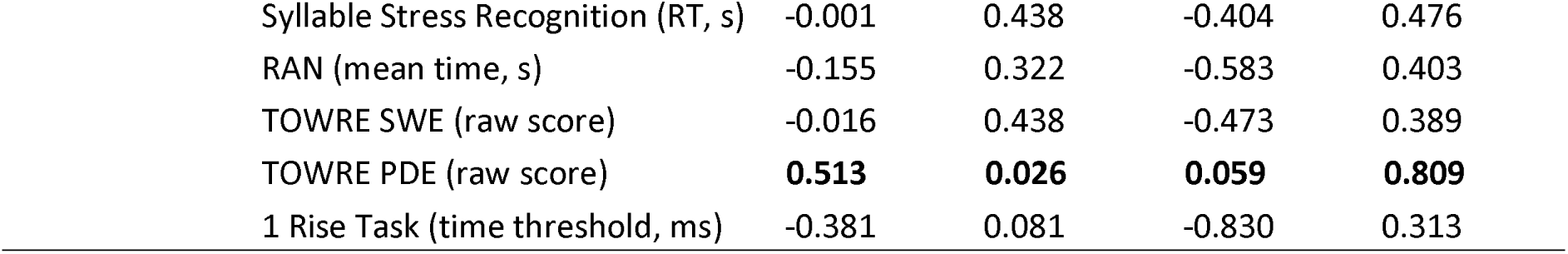
Bend correlations between BCI score and behavioral tests by group, more stringent offline preprocessing only. One-tailed p-values were FDR-corrected within each analysis set. CI95% indicates the 95% bootstrapped confidence interval for the bend correlation coefficient.

#### 3.3.2 All BCI Learners: Bend Correlations with Improvements in Behavioural Tasks

Finally, given that many control participants also demonstrated learning of the BCI, we considered the BCI learners only as a single pooled group. Given the data shown in Table 3.4, we used the offline preprocessing measure of BCI learning only. This resulted in 16 participants, 10 with dyslexia and 6 controls. The results are shown in Table 3.5. When all BCI learners were considered as a group, the data showed significant improvements in both real word reading (r= .61, FDR-corrected p = .032, 95% CI [.21, .82], Figure 3.3(A)) after playing the BCI, and also in nonword reading (*r*= .49, FDR-corrected p = .045, 95% CI [-.02, .84], Figure 3.3(B)). The confidence interval for TOWRE SWE was entirely positive, whereas the confidence interval for TOWRE PDE was largely positive but narrowly included zero. The BCI learners also showed significant improvement in amplitude rise time discrimination, which was not predicted *a priori* (*r*= –.55, FDR-corrected p = .034, 95% CI [-.87, –.09], Figure 3.3(C)). The relevant scatter plots and regression lines are shown in Figure 3.3 (offline preprocessing). These findings suggest that the BCI was successful in improving the target skills of real word reading and phonological recoding (nonword reading) for both participants with dyslexia and for control participants, as well as improving auditory discrimination of amplitude rise times for these participants. The scatterplots demonstrate that the observed relationship holds true within the dyslexic and control populations alike. By Temporal Sampling theory, it is the poor discrimination of amplitude rise times in dyslexia that causes the phonological ‘deficit’ via impaired accuracy of speech-brain alignment and associated impairments in automatic extraction of the amplitude modulation hierarchy nested in the speech envelope. The scatterplots suggest that those who made the most substantial gains in rise time discrimination and reading were all dyslexic participants.

**Figure 3.3.**
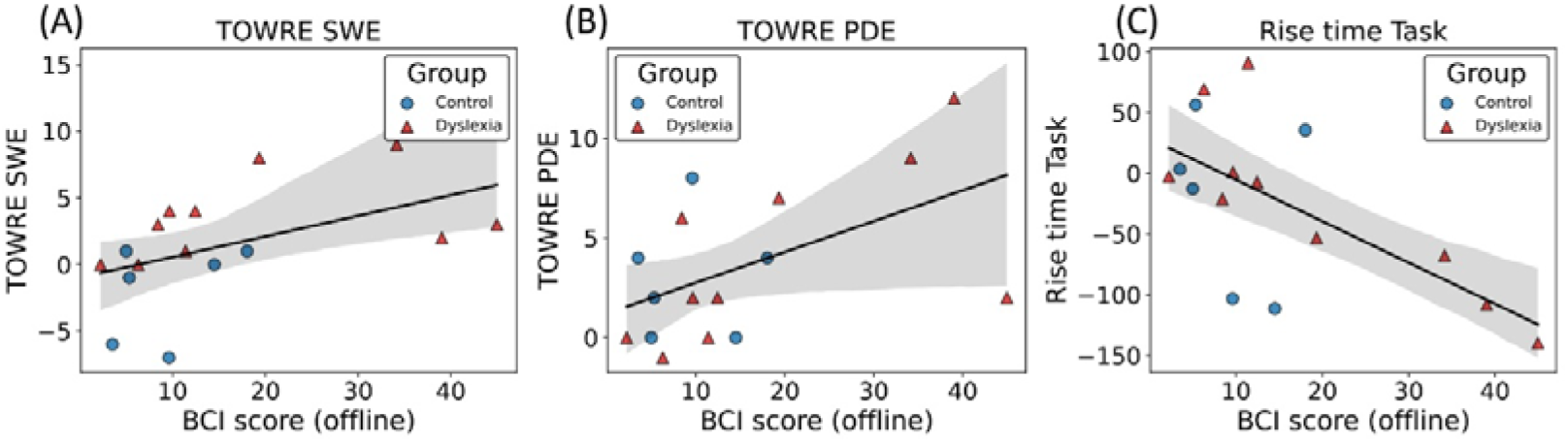
Associations between offline BCI learning scores and behavioural change scores in BCI learners. Scatter plots show the relation between offline BCI score and change in (A) TOWRE sight word efficiency, (B) TOWRE phonemic decoding efficiency, and (C) rise time threshold. Lines show the linear fit for visualisation.

**Table 3.5.** Bend correlations between BCI score and behavioral test improvements in all BCI learners (N=16), more stringent offline preprocessing protocol to remove potential artifacts, 1-tailed tests.

| BCI score<br>(t-value pre/post) | behavioral test<br>(Post - Pre) | r | pFDR | CI95% |  |
| --- | --- | --- | --- | --- | --- |
| offline<br>(all BCI learners:<br>n=16) | WRAT Arithmetic (scaled score) | -0.492 | 0.106 | -0.833 | 0.021 |
|  | WAIS Matrices (T-Score) | 0.394 | 0.131 | -0.123 | 0.804 |
|  | Phoneme Deletion (n correct) | -0.319 | 0.633 | -0.677 | 0.209 |
|  | Syllable Stress Recognition (ACC) | 0.337 | 0.126 | -0.321 | 0.792 |
|  | Syllable Stress Recognition (RT, s) | -0.174 | 0.26 | -0.648 | 0.384 |
|  | RAN (mean time, s) | -0.131 | 0.262 | -0.709 | 0.639 |
|  | TOWRE SWE (raw score) | **0.605** | **0.032** | **0.205** | **0.822** |
|  | TOWRE PDE (raw score) | **0.49** | **0.045** | **-0.018** | **0.837** |
|  | 1 Rise Task (time threshold, ms) | **-0.551** | **0.034** | **-0.871** | **-0.088** |

## Discussion

The current report suggests that a BCI aimed at normalising the low-frequency oscillatory neural patterns associated with continuous speech processing in developmental dyslexia can improve linguistic phonological processing of syllable stress patterns and phonological recoding of print to sound (nonword reading) for adults with dyslexia (Table 3.4). Further, when all BCI learners were considered together (dyslexic plus control participants), the BCI was additionally associated with significant improvements in amplitude rise time discrimination and real word reading (Table 3.5). Amplitude rise time is an important acoustic cue used for automatic oscillatory phase-resetting during speech-brain alignment (Doelling et al., 2014), and is poorly discriminated by participants with dyslexia. The gains found in syllable stress processing, real word and nonword reading for successful BCI learners are in accord with Temporal Sampling theory and appear very promising. The gains found in amplitude rise time discrimination were not predicted but are theoretically interesting, as well as being indicative of the promise of the current BCI approach for improving phonological development in children with dyslexia.

Temporal Sampling theory is based on atypical encoding of the low-frequency envelope information < 10 Hz thought to govern prosodic perception (Goswami, 2011). The improvements in syllable stress processing following BCI training for the participants with dyslexia (Table 3.4; offline preprocessing protocol only) are exciting, and on the basis of related Temporal Sampling-driven speech modelling work, should have positive effects on phonological processing at all levels of the linguistic hierarchy (Goswami, 2018). These speech modelling studies indicated that sensory discrimination of the phase relations between amplitude modulations at the delta (0.5 – 4 Hz) and theta (4 – 8 Hz) rates determines whether adult listeners perceive a strong versus a weak syllable (Leong et al., 2014). Here, when adult dyslexic participants were successful in reducing their theta-delta ratio during natural speech listening, the accuracy of their syllable stress judgements improved. The prior modelling also indicated that when one amplitude modulation cycle at each level of the amplitude modulation hierarchy is assumed to match one linguistic unit, the model extracts syllables, stressed syllables and onset-rime units with >90% accuracy for rhythmic child-directed speech (Leong & Goswami, 2015). Accordingly, improving prosodic processing in dyslexia should also improve syllabic and onset-rime processing. No associations were found for phoneme-level linguistic tasks in any analyses, however this could reflect ceiling effects on the phoneme deletion task that was selected for this study. It may also be necessary to focus a BCI on faster oscillatory responses in order to change phoneme processing in dyslexia (Marchesotti et al., 2020).

For both adults with dyslexia (Table 3.4; offline preprocessing protocol only) and all pooled BCI learners (Table 3.5; offline preprocessing protocol), BCI scores were significantly associated with enhanced nonword reading. When all the BCI learners were pooled together, BCI scores also showed a significant correlation with real word reading. These effects appear highly promising therapeutically, as the eventual target of the BCI is children with dyslexia. The compromised reading skills that ensue from the phonological processing difficulties in developmental dyslexia cause affected children to experience repeated failure in comparison to their peers, and they begin to avoid reading (Snowling et al., 2007). Playing a BCI that changes neural responses during natural speech listening before schooling begins could thus prevent such repeated failure. The BCI developed here was based on story listening, not training in print, yet it improved participants’ reading skills. As impaired nonword reading in particular is a hallmark of childhood dyslexia across languages, further development of the current BCI for children may offer significant educational benefits.

By Temporal Sampling theory, both impaired amplitude rise time discrimination and associated impaired neural encoding of low-frequency speech information compromise the efficient development of a phonological lexicon in dyslexic individuals. Indeed, experimental work with dyslexic adults has demonstrated such a relationship regarding impaired amplitude rise time discrimination and impaired speech encoding (Lizarazu et al., 2021). Meanwhile, a series of behavioural studies with dyslexic children across languages (summarized in Goswami, 2015) demonstrate that impaired amplitude rise time discrimination is significantly related to impairments in phonological awareness at many linguistic levels (e.g., tone awareness in Chinese, syllable awareness in Spanish). Here, when all BCI learners were pooled together (offline preprocessing protocol), the BCI also resulted in improved amplitude rise time discrimination. Although this was not predicted, it is theoretically interesting. Accordingly, learning the current BCI in childhood could also enhance ART discrimination, with positive effects on children’s developmental trajectories for phonological development.

To our knowledge, this is the first BCI for dyslexia that targets the pre-reading ‘phonological deficit’. The fact that it improved reading performance in this sample of adults would be expected theoretically. By the Temporal Sampling theory of dyslexia, the auditory organization of speech information by a child (assigning acoustic elements of speech perception to the groupings comprising words in a particular language) is impaired at the prosodic level. This leads to developmental differences in the accuracy of stored phonological representations for words at the level of syllable stress patterning. However, as the prosodic or rhythmic level is the foundational perceptual level regarding the rest of the linguistic hierarchy (syllables, onset-rimes and phonemes, see Leong & Goswami, 2015), these inaccurate prosodic representations affect all levels of phonological representation for affected children. This difference in phonological representations means that when print is encountered and visual codes for representing spoken language are acquired (culturally-specific codes that are taught and learned using symbol-sound correspondences), the dyslexic child is at a disadvantage from the outset. If the current BCI could be applied with children prior to learning to read, this disadvantage could potentially be eliminated before school entry. Indeed, brain imaging studies across languages show that visual symbol learning, whether of the alphabet or of characters such as Kanji, is linked to sound from the very beginning of acquiring reading (Blau et al., 2010; Froyen et al., 2009; Maurer et al., 2005, 2011; Yang et al., 2020). Accordingly, by targeting neural features of the dyslexic brain’s response to acoustic linguistic input (natural speech) before reading instruction commences, the current BCI may be able to facilitate visual symbol learning in any language.

The current study has a number of limitations. Firstly, the training sessions were given over a relatively short period of time, and half of the participants did not succeed in learning the BCI when assessed by the more stringent offline preprocessing protocol (10/20 dyslexics, 6/12 controls). As both control participants and dyslexic participants failed to learn the BCI, an explanation based on reduced listening attention in dyslexia appears unlikely. One plausible explanation could be insufficient gaming experience, accordingly a longer gaming period may be beneficial in studies which involve child participants. Secondly, we do not have information about how the participants who learned the BCI were successful. Although we asked all the participants about their listening strategies after the experiment, most were unable to report a cognitive strategy. Some participants were able to indicate chosen strategies which appeared congruent with our design intentions, for example “letting your brain flow up and down with the syllables in speech in a new way”, and “I always tried to make my mind move up and down with the flow of the speech”. However, others voiced strategies that were unlikely to have been relevant to speech processing, such as “I tried to visualize the story in my head as I listened” and “I imagined what would happen after the part I was currently listening to”. As biofeedback is operant learning, it is not necessarily surprising that participants had no conscious awareness of how they were changing their neural responses during speech listening.

A third limitation is that the sample size was relatively small. This was a convenience sample based on respones to our request for participants and so sample size was not based on a power calculation. However, the sample size is comparable to prior studies attempting to create BCIs for dyslexia (Günet, 2020, N=16 dyslexics for BCI, N=14 dyslexic controls); Christodoulides et al., 2022, N=14 dyslexics, N=12 controls). Furthermore, EEG data are prone to exhibiting highly variable day-to-day variations. To mitigate this problem, baseline data was collected before each BCI run and these recorded EEG patterns were used to define the upper and lower limits of the spaceship on the screen on each run. Because the feedback scale was recalibrated before each run, the BCI score was not directly comparable across runs or sessions, limiting the computation of a continuous online measure of BCI learning. An additional limitation is that a single story (Winnie the Pooh) was used throughout the whole training protocol. The story was written in simple child language and did not require any effort regarding comprehension. While this decision was helpful in allowing direct comparison of performance across sessions, it also made the protocol quite tedious, which may have led to de-motivation (Alkhadim, 2022) or may have negatively affected the dyslexic participant with ADHD (this participant had a significant t-statistic in the online learning protocol but not in the offline protocol). However, demotivation should apply equally to all participants. Following a reviewer suggestion, we also computed alpha and beta power across the study, to see whether these potential indices of attention varied by group. Alpha and beta power in sessions 1 and 16 was equivalent by group and was also equivalent for learners and non-learners. Nevertheless, given that the intended target population is young children, it would be best if future work could devise an operant learning protocol that can utilise many different stories to ensure motivation and attentional engagement.

Finally, while the provided instructions were quite clear regarding the gaming objective (i.e. making the spaceship go upwards as consistently as possible on the screen while listening to the words in the story carefully), the instructions were also kept purposefully vague so that participants could decide by themselves which strategy to employ. Regarding the target audience of children, it is possible that more direct instructions about potential listening strategies could be useful, for example instructing participants to try to listen to syllable patterning and speech rhythm. It is also possible that some participants used subvocalization, which would change the task from one of passive speech processing to active speech production, thereby introducing a potential muscle artifact. However, no participants were observed to use subvocalization. Further, due to the offline filtering applied, any high frequency signal linked to muscle artifacts was unlikely to affect spaceship position.

In conclusion, the exploratory data presented here suggest that a simple and engaging BCI for improving phonological processing during natural speech listening can be created using EEG data informed by the Temporal Sampling theory of developmental dyslexia. Participants who learned the BCI showed improved processing of syllable stress patterns in words, improved phonological recoding skills (nonword reading), improved word reading skills, and improved amplitude rise time discrimination. These improvements occurred even though no direct training of phonology, word or nonword reading nor auditory discrimination occurred during the study. This is interesting theoretically, as it suggests that the therapeutic benefits resulted from improving the neural theta-delta ratio during natural speech listening. Therapeutic interventions which filter speech to enhance amplitude rise times have also been shown to improve speech processing in participants with dyslexia via changing the theta-delta ratio (Mandke et al., 2023). If such a filter could be integrated with a form of hearing aid, this could in principle provide a convergent therapy for dyslexia to a BCI. Accordingly, further investigation of neural speech processing in dyslexia guided by Temporal Sampling theory be beneficial, and may identify other, possibly more effective, neural targets for BCI development.

## ACKNOWLEDGEMENTS

The authors would like to thank all the participants who volunteered for the study. This research was funded by a donation to U.G. from the Yidan Prize Foundation. The sponsor played no role in the study design, data interpretation, nor writing of the report.

## DATA AND CODE AVAILABILITY

Data and code will be made available on request.

## Abbreviations

ACC: Accuracy
ADHD: Attention deficit hyperactivity disorder
ART: Amplitude rise time
ASR: Artifact Subspace Reconstruction
BCI: Brain-computer interface
BF10: Bayes factor in favour of the alternative hypothesis
CI: Confidence interval
CTRL: Typically-developing control group
DYS: Dyslexia group
EEG: Electroencephalography
EI: Efficiency index
EMG: Electromyography
EOG: Electrooculography
FDR: False discovery rate
HCPC: Health and Care Professions Council
ICA: Independent Component Analysis
PDE: Phonemic Decoding Efficiency
PSD: Power spectral density
RAN: Rapid Automatized Naming
RT: Reaction time / response time
SD or S.D.: Standard deviation
SpLD: Specific Learning Difficulties
SWE: Sight Word Efficiency
TOWRE: Test of Word Reading Efficiency
WAIS: Wechsler Intelligence Scale for Adults
WRAT: Wide Range Achievement Test

